# Context-dependent regulation of MGMT expression by IRE1 signaling and its relationship to etoposide-associated DNA damage in MDA-MB-231 cells

**DOI:** 10.64898/2026.11.24.768915

**Authors:** Kevin Y. Chen, Elaina Gouin, Ava Erickson, Samantha Zaloudek, Sean Foster, Caleb Sandum, S. Patrick Walton, Christina Chan

## Abstract

Inositol-requiring enzyme 1α (IRE1) links endoplasmic reticulum stress to adaptive gene regulation, but its relationship with O⁶-methylguanine-DNA methyltransferase (MGMT) expression in breast cancer remains unclear. We investigated IRE1-associated MGMT regulation and etoposide-associated DNA damage in IRE1-proficient and IRE1-deficient MDA-MB-231 cells. IRE1 deficiency reduced basal MGMT mRNA and protein levels. MGMT overexpression reduced etoposide-associated comet tail moment in both genotypes but did not eliminate the genotype-associated difference, suggesting contributions from additional IRE1-dependent mechanisms. IRE1 re-expression increased MGMT mRNA under vehicle and decitabine treatment, whereas X-box binding protein 1s (XBP1s) overexpression increased MGMT mRNA only under decitabine treatment. Neither intervention significantly increased MGMT protein level relative to its corresponding vector control. Chromatin immunoprecipitation analysis showed XBP1s association with the assayed MGMT promoter region under decitabine treatment. Combined palmitate and etoposide exposure reduced MGMT mRNA in IRE1-proficient but not IRE1-deficient cells, and this response persisted following XBP1 depletion. MGMT mRNA association with IRE1-containing complexes and a predicted RNA recognition element supported potential regulated IRE1-dependent decay-like regulation, without establishing direct cleavage. Together, these findings support context-dependent IRE1-associated regulation of MGMT, linking maintenance of basal expression with transcript loss during combined metabolic and genotoxic stress in MDA-MB-231.

## Introduction

The regulation of DNA repair genes can influence both genome maintenance and cancer-cell responses to treatment (ref. 1-3). O^6^-methylguanine-DNA methyltransferase (MGMT) is a key DNA repair protein that protects the genome from alkylation damage by removing alkyl groups from the O^6^ position of guanine. This repair is through irreversible transfer of the alkyl group to an active-site cysteine residue on MGMT, thereby restoring the unmodified guanine base while simultaneously inactivating the MGMT enzyme (ref. 4-5). Although this canonical lesion-reversal activity is distinct from the direct repair of DNA double-strand breaks (DSBs), MGMT suppression has been reported to impair homologous-recombination-associated responses and enhance cisplatin-induced DSBs in nasopharyngeal carcinoma models (ref. 6). Because MGMT expression and promoter methylation can influence cellular responses to DNA-damaging therapies, they have been widely investigated as potential determinants of treatment response. However, their clinical significance is context dependent and should be evaluated within specific cancer types and treatment regimens.

Many cancers experience chronic endoplasmic reticulum (ER) stress and activate the unfolded protein response (UPR). Increasing evidence suggests there is crosstalk between the ER proteostasis and DNA damage repair pathways (ref. 7). Inositol-requiring enzyme 1α (IRE1), one of the three canonical ER stress sensor proteins of the UPR, offers a potential connection between cellular stress adaptation and the regulation of DNA-damage responses. IRE1 RNase signaling has two major outcomes: it generates the spliced, transcriptionally active form of X-box binding protein 1 (XBP1s) through noncanonical XBP1 mRNA splicing and reduces selected transcripts through regulated IRE1-dependent decay (RIDD) (ref. 8-10). RNA-substrate specificity for RIDD depends on the activation state of IRE1: dimeric complexes favor cleavage at defined endomotifs, whereas higher-order phosphorylated assemblies can promote more broadly permissive RNA decay (ref. 9). Thus, a putative recognition motif alone is insufficient to designate a transcript as an IRE1 decay target (ref. 9). IRE1 signaling also contributes to breast cancer phenotypes through several experimentally demonstrated mechanisms. XBP1 was shown to support survival of triple-negative breast cancer through cooperation with the HIF1α pathway (ref. 11). In contrast, pharmacological inhibition of IRE1 RNase activity reduced production of tumor-promoting cytokines and enhanced paclitaxel-mediated tumor suppression in preclinical triple-negative breast cancer models (ref. 12). These findings establish the importance of IRE1-XBP1 signaling in breast cancer adaptation and stress tolerance. They provide evidence that IRE1-XBP1 signaling contributes to tumor cell survival; however, these findings do not, by themselves, demonstrate that MGMT is regulated by IRE1-XBP1 pathway, either directly or through downstream intermediates.

In glioblastoma, elevated MGMT expression contributes to temozolomide (TMZ) resistance. In this context, XBP1s transcriptionally activated α-synuclein, whose expression reduced MGMT levels and partially restored TMZ sensitivity in TMZ-resistant U87 cells (ref. 13). These findings do not establish direct XBP1s regulation of the MGMT promoter, nor does it demonstrate that MGMT mRNA is an IRE1 cleavage substrate (ref. 13). The present study investigated the relationship between IRE1 signaling, MGMT expression, and etoposide-associated DNA damage in MDA-MB-231 cells. Using IRE1-proficient and IRE1-deficient cells, MGMT overexpression, IRE1 re-expression, XBP1s overexpression, decitabine treatment, and chromatin- and RNA-association analyses, we examined potentially distinct transcriptional and post-transcriptional mechanisms of MGMT regulation. Previously our laboratory showed that palmitate engages IRE1-XBP1 signaling in breast cancer cells (ref. 14), consistent with evidence that membrane lipid saturation can activate IRE1 through its transmembrane domain (ref. 15-16), therefore we investigated the regulation of MGMT under combined palmitate and etoposide exposure mediated by IRE1-dependent RIDD-like regulation.

## Results

### IRE1 deficiency reduces basal MGMT expression and limits protection associated with MGMT overexpression

To investigate the relationship between IRE1 and MGMT in breast cancer cells, we first compared basal MGMT expression between wild-type (WT) and IRE1-deficient (IRE1^-/-^) MDA-MB-231 cells. MGMT mRNA level was significantly reduced in IRE1^-/-^ cells relative to WT cells (**Fig. 1a**), accompanied by lower MGMT protein level (**Fig. 1b**). These findings support a role for IRE1 signaling in controlling MGMT expression.

**Figure 1.**
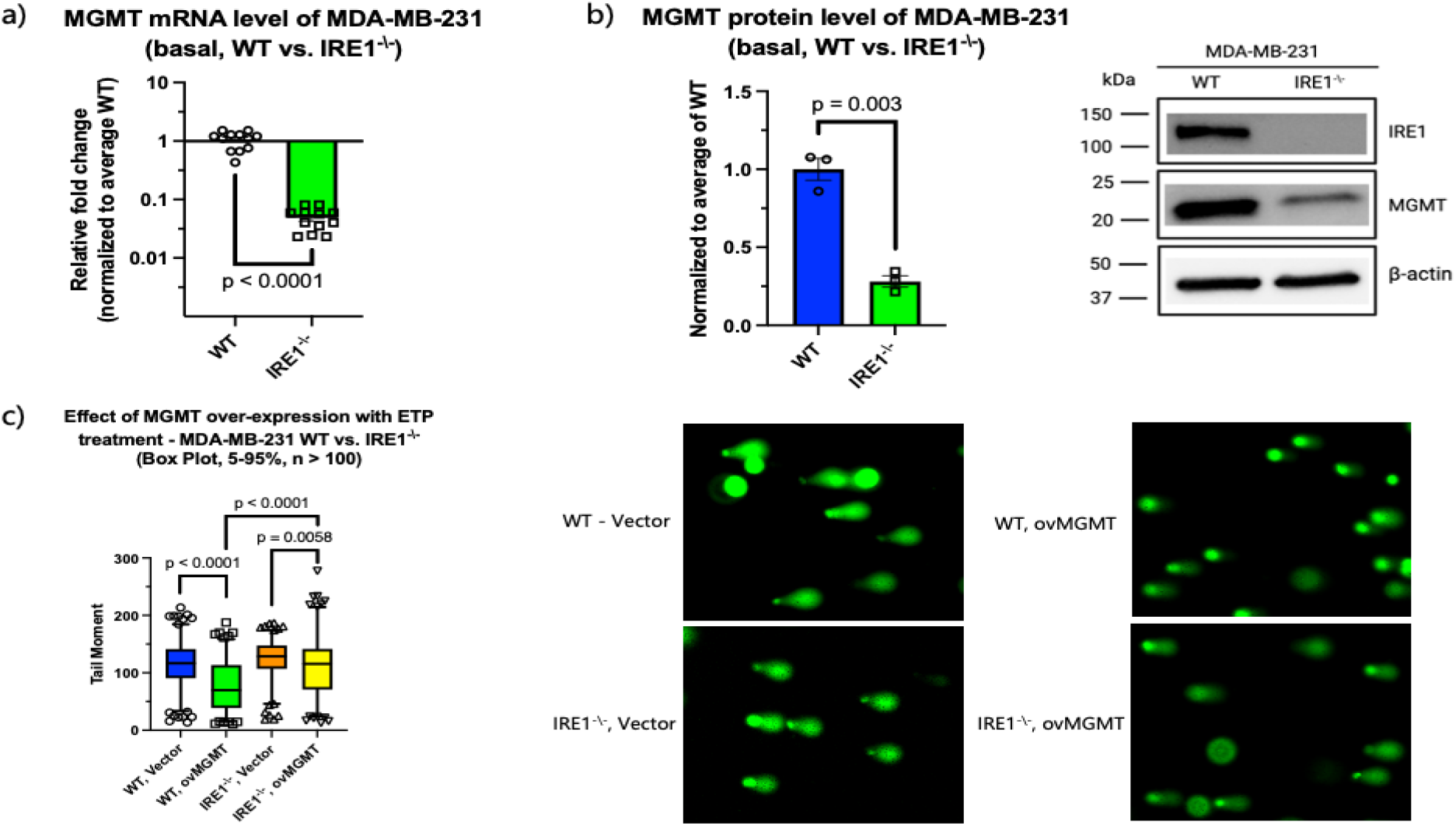
Relationship between IRE1, MGMT expression, and etoposide-associated DNA damage in MDA-MB-231 cells. (a) Quantification of MGMT mRNA via qRT-PCR analysis between IRE1 WT and IRE1^-/-^ cells. (b) Western blot analysis of MGMT and IRE1 protein expression between IRE1 WT and IRE1^-/-^ cells. Both mRNA and protein levels were normalized to the average of WT (y-axis). Data are mean ± SEM from n = 12 independent biological replicates for qRT-PCR and n = 3 independent biological replicates for western blot. (c) Comet assay of MDA-MB-231 WT and IRE1^-/-^ cells overexpressing MGMT (ovMGMT) exposed to ETP. Each condition was measured in 3 independent runs with > 50 cells measured per run. Plots show median (horizontal line), 25th and 75th percentiles (bottom and top of the box, respectively), and 5th and 95th percentiles (bottom and top error bars). A representative image for each condition is also shown.

We next examined whether MGMT overexpression reduced etoposide (ETP)-associated DNA damage in the presence or absence of IRE1. Given that MGMT is lowly expressed in MDA-MB-231 cells (ref. 18), MGMT was overexpressed in WT and IRE1^-/-^ cells, and DNA damage following ETP treatment was assessed using the comet assay. MGMT overexpression significantly reduced comet tail moment in both genotypes relative to their corresponding empty-vector controls (**Fig. 1c**, green vs. blue in WT and yellow vs. orange in IRE1^-/-^ cells). However, MGMT-overexpressing IRE1^-/-^ cells retained significantly higher tail moments than MGMT-overexpressing WT cells (**Fig. 1c**, yellow vs. green). Thus, although MGMT overexpression reduced ETP-associated DNA damage in both backgrounds, it did not eliminate the difference between WT and IRE1^-/-^ cells. Together, these findings connect IRE1 signaling to the maintenance of basal MGMT expression and support an additional role of IRE1 in limiting DNA damage accumulation.

### IRE1 re-expression increases MGMT mRNA but not protein level under DMSO conditions in IRE1-deficient cells

Having observed reduced basal MGMT expression in IRE1-deficient cells, we next examined whether IRE1 re-expression restored MGMT level. Since MGMT expression can be regulated by epigenetic mechanisms, we determined if DNA methylation is involved in the IRE1-dependent control of MGMT expression. To investigate whether DNA methyltransferase inhibition influenced this response, we treated the cells with decitabine (5-aza-2′-deoxycytidine; DAC), a DNA methyltransferase inhibitor that promotes DNMT1 depletion (ref. 19). MDA-MB-231 IRE1^-/-^ cells were transfected with an IRE1 expression construct (reIRE1) or an empty-vector control and subsequently treated with DAC or its vehicle, DMSO. MGMT protein level did not differ significantly between reIRE1 cells and the corresponding empty-vector controls under either DMSO- or DAC-treated conditions (**Fig. 2a-b**).

**Figure 2.**
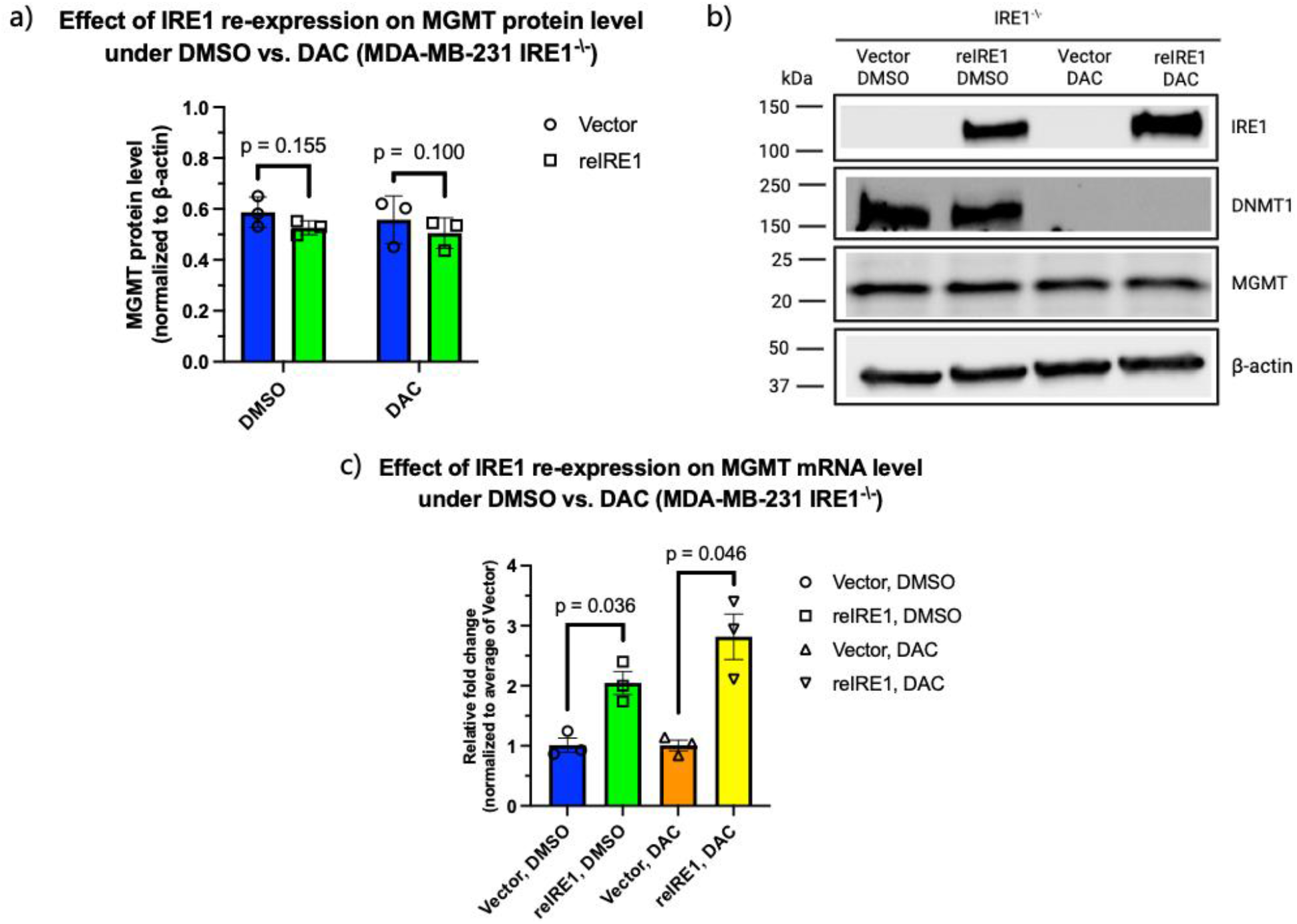
IRE1 re-expression increases MGMT mRNA level under both DMSO and DAC treatment without significantly altering MGMT protein level in MDA-MB-231 IRE1^-/-^ cells. Cells were transfected with an IRE1 expression construct (reIRE1) or an empty-vector control and treated with decitabine (DAC) or DMSO. (a) Quantification of MGMT protein level by Western blotting, normalized to β-actin. (b) Representative Western blots showing IRE1, DNMT1, MGMT, and β-actin (loading control). (c) MGMT mRNA levels measured by qRT-PCR, normalized to 18S rRNA, and expressed relative to the corresponding empty-vector control within each treatment condition (DMSO or DAC). Data are presented as mean ± SEM from n = 3 independent biological replicates.

In contrast, qRT-PCR analysis showed that IRE1 re-expression significantly increased MGMT mRNA levels relative to the corresponding vector controls under both DMSO and DAC treatment (**Fig. 2c**). For these comparisons, MGMT mRNA levels were normalized to the vector control within each treatment condition. These increases were consistent with the lower basal MGMT mRNA levels observed in IRE1-deficient cells relative to WT cells (**Fig. 1a**). Thus, IRE1 re-expression increased MGMT transcript levels under both treatment conditions without a corresponding detectable increase in MGMT protein (**Fig. 2a-b**). Because IRE1 re-expression alone does not necessarily restore activation of its downstream XBP1 pathway, we next examined whether expression of the spliced transcription factor XBP1s is sufficient to regulate MGMT expression independently of endogenous IRE1.

### XBP1s overexpression increases MGMT mRNA under DAC treatment and is associated with MGMT-promoter occupancy

To examine the contribution of XBP1s independently of endogenous IRE1, we overexpressed XBP1s (ovXBP1s) in MDA-MB-231 IRE1^-/-^ cells and measured MGMT expression following DMSO or DAC treatment. MGMT protein level, normalized to β-actin, did not differ significantly between ovXBP1s cells and the corresponding empty-vector controls under either treatment condition (**Fig. 3a–b**). In contrast, qRT-PCR analysis showed that XBP1s overexpression significantly increased MGMT mRNA levels relative to the corresponding vector control under DAC treatment, whereas no significant difference was detected under DMSO treatment (**Fig. 3c**). MGMT mRNA levels were normalized to 18S rRNA and expressed relative to the empty-vector control within each treatment condition. XBP1s overexpression increased MGMT transcript levels under DAC conditions without a corresponding detectable increase in MGMT protein.

**Figure 3.**
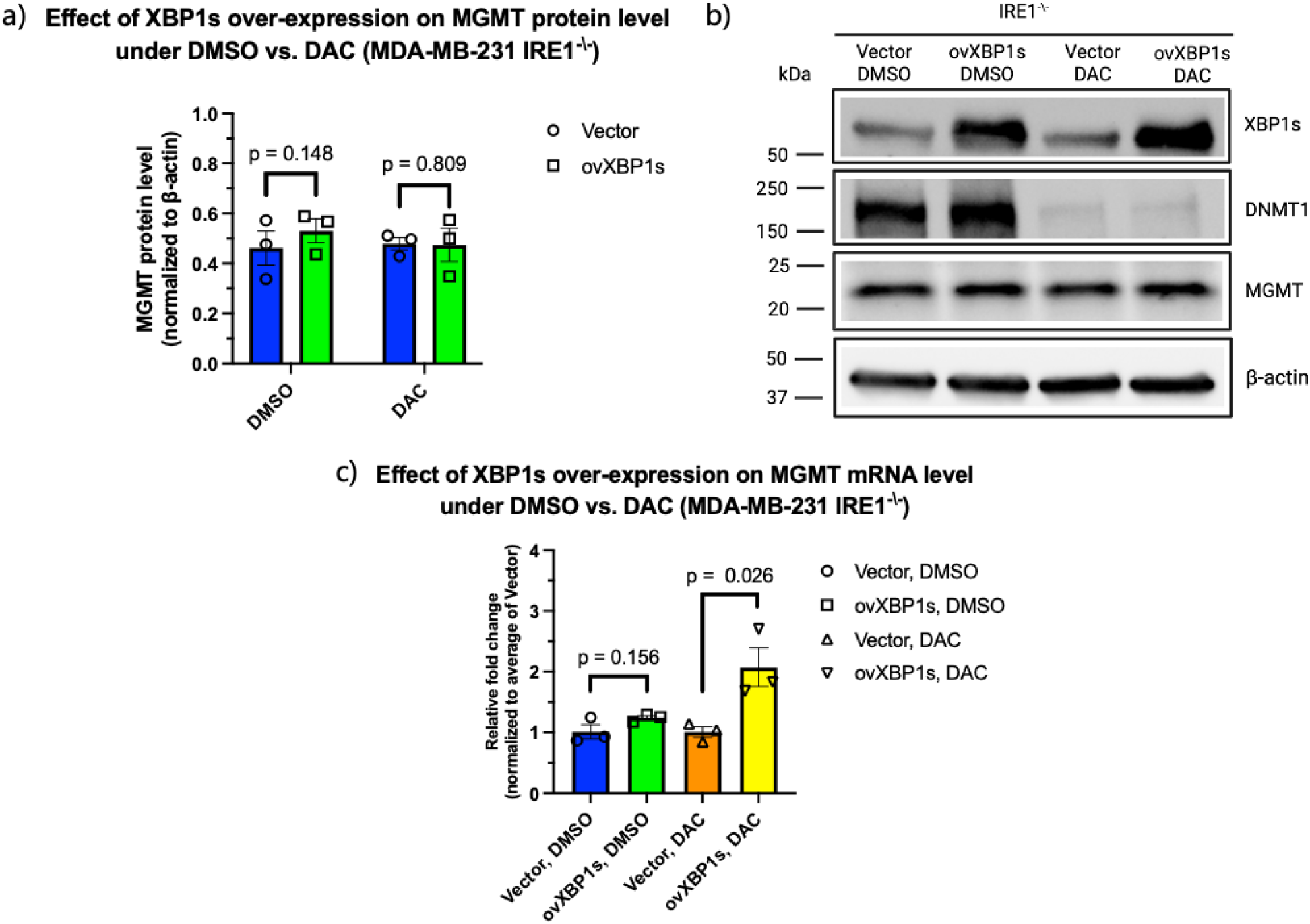
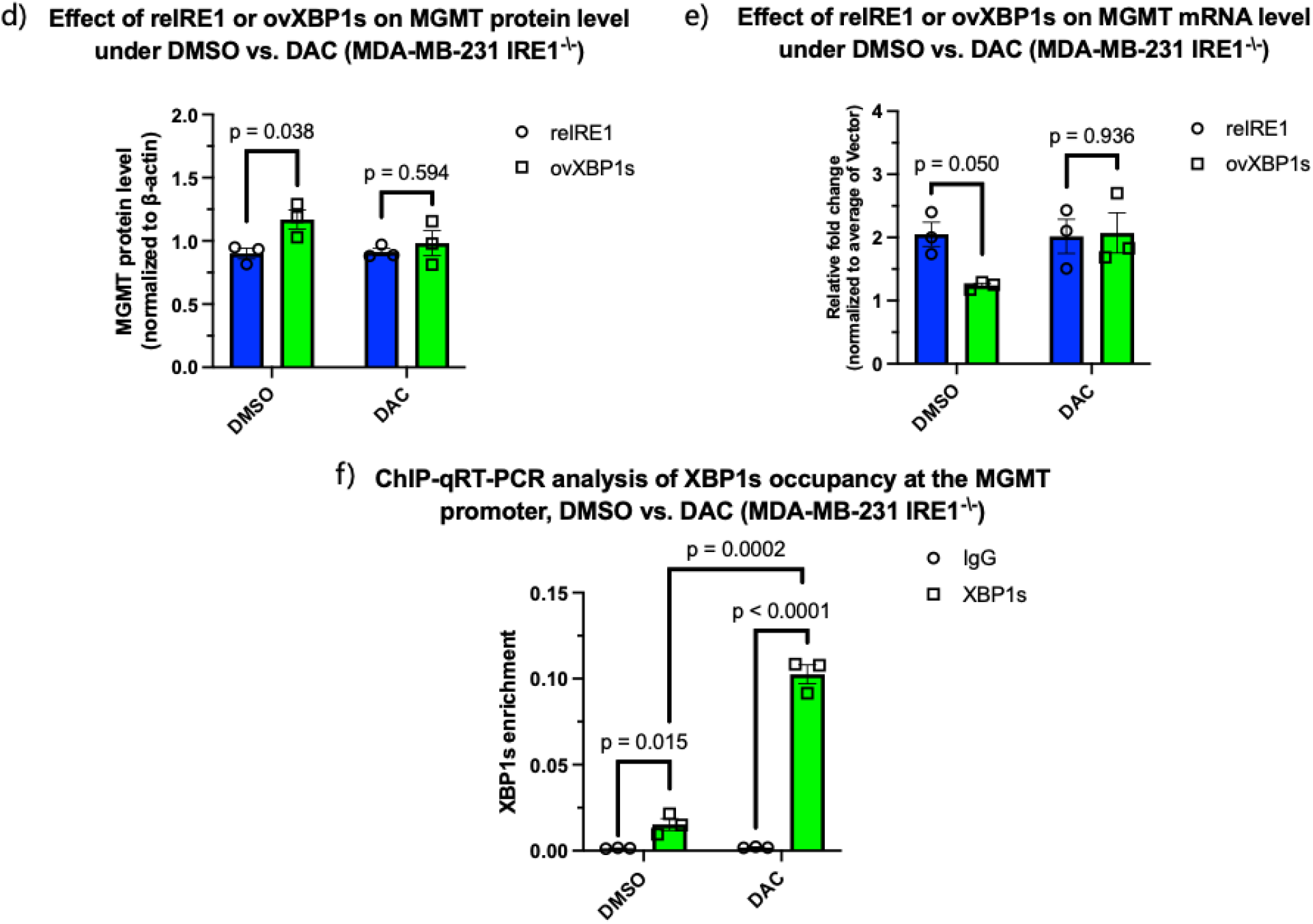
MGMT expression following XBP1s overexpression, comparison with IRE1 re-expression, and XBP1s association with the MGMT promoter. Cells were transfected with an XBP1s expression construct (ovXBP1s) or an empty-vector control and treated with DAC or DMSO. (a) Quantification of MGMT protein level by Western blotting, normalized to β-actin. (b) Representative Western blots showing XBP1s, DNMT1, MGMT, and β-actin, which served as the loading control. (c) MGMT mRNA levels measured by qRT-PCR, normalized to 18S rRNA, and expressed relative to the corresponding empty-vector control within each treatment condition (DMSO or DAC). Comparison of MGMT protein (d) and mRNA (e) levels between reIRE1 and ovXBP1s under DMSO- or DAC-treated conditions. Data were derived from the experiments presented in Fig. 2 and 3. Data are presented as mean ± SEM from n = 3 independent biological replicates. (f) ChIP-qPCR analysis of XBP1s association with the MGMT promoter in XBP1s-overexpressing MDA-MB-231 IRE1^-/-^ cells treated with DMSO or DAC. IgG served as the negative immunoprecipitation control. Samples from three independent biological experiments were pooled within each treatment condition, yielding one pooled sample per condition. Each pooled sample was analyzed in technical triplicate. Data are presented as mean ± SEM of the technical measurements; error bars reflect technical variability, not variability among independent biological experiments.

We next compared MGMT expression following XBP1s overexpression and IRE1 re-expression. At the protein level, β-actin-normalized MGMT level was modestly but significantly higher in ovXBP1s cells than in reIRE1 cells under DMSO treatment (p = 0.038), whereas no significant difference was detected under DAC treatment (**Fig. 3d**). Despite this difference between the two expression conditions under DMSO, neither intervention significantly altered MGMT protein level relative to its corresponding empty-vector control under either treatment condition (**Fig. 2a–b**; **Fig. 3a–b**).

At the mRNA level, the comparison was based on 18S rRNA-normalized MGMT expression relative to each construct’s corresponding empty-vector control within each treatment condition. Under DMSO, this vector-normalized response was lower following XBP1s overexpression than following IRE1 re-expression (p = 0.05, **Fig. 3e**, DMSO pair). No significant difference was detected under DAC treatment (**Fig. 3e**, DAC pair). Relative to their respective vector controls, IRE1 re-expression significantly increased MGMT mRNA under both treatments, whereas a significant increase following XBP1s overexpression was detected only under DAC. Together, these findings indicate that the two interventions produced different patterns of MGMT transcript response relative to their controls, without corresponding significant increases in MGMT protein level.

Because XBP1s overexpression increased MGMT mRNA levels under DAC treatment (**Fig. 3c**), we next examined whether XBP1s is associated with the MGMT promoter. ChIP-qPCR analysis was performed by Creative BioMart using XBP1s-overexpressing MDA-MB-231 IRE1^-/-^ cells treated with DMSO or DAC. Under DAC treatment, enrichment of the assayed MGMT-promoter region was greater in XBP1s immunoprecipitates than in IgG controls. Enrichment in XBP1s immunoprecipitates was much greater following DAC treatment than DMSO treatment (**Fig. 3f**). Together with the increased MGMT mRNA levels observed following XBP1s overexpression under DAC treatment, these findings suggest a potential transcriptional contribution of XBP1s to MGMT regulation in this setting.

### Palmitate and etoposide co-treatment reduces MGMT mRNA in a pattern consistent with RIDD activity

IRE1 RNase signaling includes both XBP1 mRNA splicing, which generates the transcription factor XBP1s, and regulated IRE1-dependent decay (RIDD) of other RNA substrates (ref. 9-10). To assess whether RIDD might contribute to MGMT regulation during combined metabolic and genotoxic stress, we depleted XBP1 via siRNA (XBP1 KD) in MDA-MB-231 IRE1 WT and IRE1^-/-^ cells and compared MGMT mRNA responses to PA+ETP versus ETP alone. This approach reduced the contribution of the canonical IRE1-XBP1 transcriptional axis while allowing us to examine whether the stress-associated response persisted after XBP1 KD and depended on IRE1 status. XBP1 KD efficiency was confirmed by qRT-PCR (**Fig. S1**).

Cells were subsequently treated with etoposide (ETP) alone or palmitate plus etoposide (PA+ETP). In WT cells, PA+ETP significantly reduced MGMT mRNA levels relative to ETP alone. The established RIDD-associated transcripts SPARC and BLOC1S1 were similarly decreased under the same treatment condition (**Fig. 4a**). These decreases persisted following XBP1 KD, showing that stress-associated transcript loss remained detectable despite reduced XBP1 expression. In contrast, PA+ETP did not significantly reduce MGMT, SPARC, or BLOC1S1 mRNA in IRE1^-/-^ cells, regardless of XBP1 KD status (**Fig. 4b**). Together, the reduction in MGMT mRNA alongside established RIDD-associated transcripts in WT but not IRE1^-/-^ cells is consistent with a contribution of IRE1-mediated RNA decay to this response.

**Figure 4.**
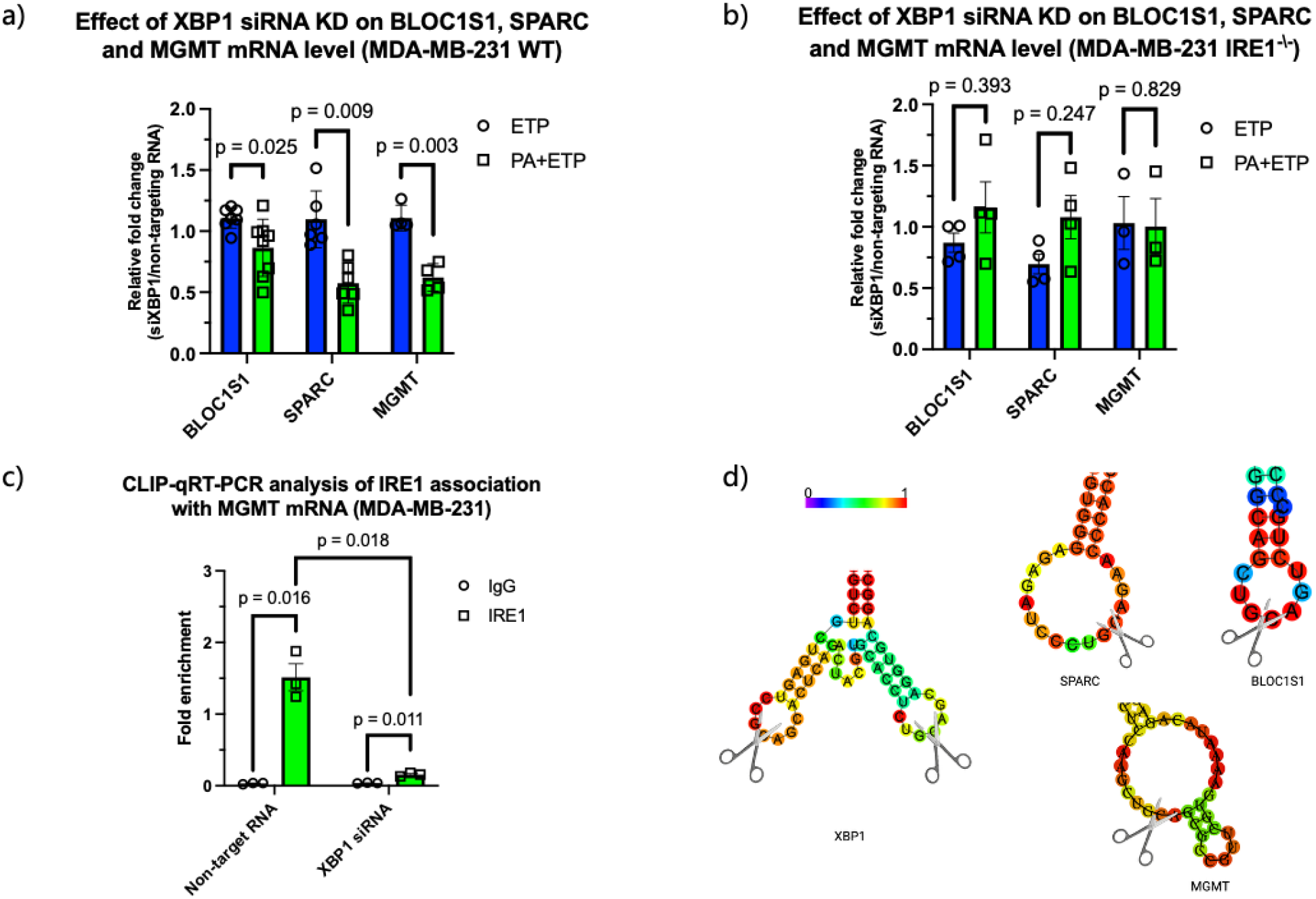
PA and ETP co-treatment promote IRE1-dependent, XBP1-independent downregulation of MGMT mRNA consistent with RIDD-like regulation. qRT-PCR measures mRNA of BLOC1S1, SPARC, and MGMT under PA+ETP co-treatment with XBP1 KD and NT between (a) WT and and (b) IRE1^-/-^. Data are mean ± SEM from n = 8, 6, 4 independent biological replicates for BLOC1S1, SPARC and MGMT, respectively in WT. For IRE1^-/-^, data are mean ± SEM from n = 4, 4, 3 independent biological replicates for BLOC1S1, SPARC and MGMT, respectively. (c) IRE1 CLIP-qPCR analysis of MGMT mRNA association with IRE1-containing complexes in PA+ETP-treated WT cells with XBP1 KD or non-targeting control. Samples from three independent biological experiments were pooled within each condition, yielding one pooled sample per condition. Each pooled sample was analyzed in technical triplicate. Data are presented as mean ± SEM of the technical measurements; error bars reflect technical variability rather than variability among independent biological experiments. (d) *In silico* RNA secondary-structure predictions for spliced XBP1, SPARC, BLOC1S1, and MGMT transcripts generated using RNAfold. XBP1, SPARC and BLOC1S1 are established IRE1-associated transcripts, and were included as positive controls. Predicted stem-loop structures containing the putative IRE1 RNase consensus endomotif, CUGCAG, are shown. Scissors indicate predicted IRE1 cleavage sites within the CUGCAG-containing stem-loop structures. Colors indicate base-pairing probabilities from 0 (low probability) to 1 (high probability).

In an independent basal-condition experiment, pharmacologic inhibition of IRE1 RNase activity with 4μ8C increased MGMT mRNA level relative to DMSO vehicle control after 6h of treatment; however, this effect was not observed after 48h (**Fig. S2**). Although this experiment did not include PA+ETP treatment or parallel measurement of established RIDD substrates, nonetheless the result is consistent with IRE1 RNase activity contributing to context-dependent regulation of MGMT mRNA levels (**Fig. 4**).

To examine whether MGMT mRNA associates with IRE1 under these conditions, IRE1 CLIP-qPCR analysis was performed by Creative BioMart. MGMT mRNA was detected in IRE1-associated RNA complexes following PA+ETP treatment, with reduced enrichment observed after XBP1 depletion (**Fig. 4c**). *In silico* analysis also identified a candidate IRE1 recognition sequence within a predicted stem-loop-like structure in the MGMT transcript (**Fig. 4d**). Together, the IRE1 dependence of PA+ETP-associated MGMT mRNA loss, the CLIP evidence of MGMT mRNA association with IRE1-containing complexes, and the predicted sequence-structure element are consistent with context-specific, RIDD-like regulation. These findings support MGMT as a candidate IRE1-regulated transcript and potential RIDD target under combined PA and ETP stress.

## Discussion

This study identified that MGMT regulation is a context-dependent component of IRE1 signaling in MDA-MB-231 cells. This relationship involves both maintenance of basal MGMT expression and its downregulation during combined metabolic and genotoxic stress, rather than a uniformly positive IRE1-XBP1s-MGMT regulation pathway. This distinction extends the established connection between ER proteostasis and the DNA-damage response (ref. 7). Previous studies have linked IRE1-XBP1 signaling to triple-negative breast cancer adaptation through HIF1α-associated transcription and regulation of the tumor-cell secretome (refs. 11-12). Our findings expand this framework by implicating the regulation of a DNA repair factor. The association between MGMT abundance and ETP-associated DNA damage is particularly noteworthy because MGMT is traditionally associated with reversal of O⁶-alkylguanine lesions rather than directly repairing ETP-induced DNA breaks. In the same breast cancer cell line, Lin et al. (2020) reported that MGMT overexpression suppressed proliferation and invasive capacity, accompanied by increased PTEN expression and reduced AKT phosphorylation (ref. 18). Evidence that MGMT suppression attenuates homologous-recombination-associated responses in cisplatin-treated nasopharyngeal carcinoma cells further links MGMT to functions beyond its canonical lesion-reversal activity (ref. 6). Together, these studies provide complementary contexts for interpreting the MGMT-associated protection observed here, including effects on cellular signaling pathways and DNA repair-associated processes. However, the persistence of the genotype-associated difference despite MGMT overexpression suggests that additional IRE1-dependent mechanisms contribute to ETP-associated DNA-damage accumulation (**Fig. 1c**). Thus, the IRE1-MGMT relationship identified in this study likely operates within a broader stress-response network.

The regulatory relationship also appears more complex than a simple transcriptional relay from IRE1 through XBP1s to MGMT. The differing responses to IRE1 re-expression and XBP1s overexpression suggest that restoring the upstream sensor and expression of its downstream transcription factor are not functionally equivalent (**Fig. 2c**; **Fig. 3c,e**). This distinction is consistent with the multiple RNA-processing outputs of IRE1 and their dependence on its activation state (ref. 9). The response observed with XBP1s under DAC may reflect a permissive regulatory context, involving promoter accessibility or cooperating factors. XBP1s association with MGMT-promoter region supports a potential transcriptional contribution in this setting (**Fig. 3f**). A useful comparison is the reported XBP1s-α-synuclein-MGMT relationship in glioblastoma, in which XBP1s acted through α-synuclein in a pathway associated with reduced MGMT expression and restored TMZ sensitivity (ref. 13). Collectively, these findings caution against assigning a universal directionality to XBP1s-dependent MGMT regulation across cancer models. The departure between MGMT transcript and protein responses further suggests that RNA abundance alone does not fully capture the underlying regulatory mechanisms (**Fig. 2a-c**; **Fig. 3a-c**). Differences in translation efficiency, protein turnover, or delayed protein accumulation could contribute to this relationship (ref. 20). Additional post-transcriptional mechanisms, including altered mRNA stability and microRNA-mediated regulation, could also influence MGMT expression (ref. 21). Together, these possibilities provide a framework for interpreting the transcript-protein relationship without assigning it to a specific mechanism.

The combined-stress experiments place MGMT regulation within the broader capacity of IRE1 to integrate metabolic cues with RNA homeostasis. Our laboratory previously linked palmitate exposure to IRE1-XBP1 signaling in breast cancer cells (ref. 14), consistent with studies demonstrating that membrane lipid saturation can activate IRE1 through its transmembrane domain (refs. 15-16). The present work extends the activation of IRE1 to the regulation of a DNA repair factor during combined metabolic and genotoxic stress. The coordinate responses of MGMT and established RIDD-associated transcripts across IRE1-proficient and IRE1-deficient cells (**Fig. 4a-b**), together with the association of MGMT mRNA with IRE1-containing complexes (**Fig. 4c**) and the presence of a predicted RIDD recognition element (**Fig. 4d**), support MGMT as a candidate substrate of RIDD-like regulation (refs. 9-10). The contrasting relationship observed under basal and stress conditions need not be viewed as contradictory. Rather, they suggest that the net effect of IRE1 on MGMT reflects the balance among multiple regulatory processes operating within a given cellular context (**Fig. 1a-b**; **Fig. 4a-b**). This interpretation broadens the connection between ER stress signaling and DNA repair-factor regulation beyond transcriptional activation alone, incorporating a potential contribution from IRE-mediated RNA-decay under combined stress conditions. More broadly, these findings support a model in which MGMT expression is shaped by transcriptional and post-transcriptional mechanisms downstream of IRE1, with the relative influence of each pathway varying according to cellular state and stress condition.

Feedback mechanisms and temporal adaptation provide additional context for this regulatory balance. XBP1 deficiency has been shown to trigger feedback activation of IRE1 and enhance RIDD in mouse liver (ref. 22), offering a possible explanation for how transcript loss can persist following XBP1 depletion. However, whether a similar feedback mechanism operates in our system was not directly examined. Likewise, the time-dependent effects to 4μ8C suggest that acute inhibition of IRE1 RNase activity and prolonged pathway perturbation may have distinct consequences for MGMT regulation (**Fig. S2**). The delayed increase in IRE1 abundance following extended 4μ8C exposure may reflect cellular adaptation to sustained inhibition, although it does not, by itself, indicate restoration of RNase activity. These observations underscore the dynamic nature of IRE1 signaling network and highlight that genetic deletion, transient re-expression, XBP1 depletion, and pharmacological inhibition interrogate different aspects of pathway function. Consequently, their collective value lies not in defining a single linear regulatory mechanism, but in revealing the context dependence of IRE1-associated MGMT regulation. This distinction is particularly important when interpreting how perturbation of an ER stress sensor influences both the expression and functional contribution of a DNA repair factor. More broadly, the findings support a model in which MGMT regulation involves multiple IRE1-dependent processes whose relative contributions vary across cellular states, temporal scales, and stress conditions.

Several limitations define the mechanistic scope and generalizability of these findings. First, the experiments were performed in a single breast cancer cell-line, and chronic IRE1 deletion may have elicited adaptive changes that are not fully reversed by transient re-expression. Although MGMT overexpression demonstrates that increasing MGMT abundance can modify the ETP-associated phenotype, it does not establish that endogenous MGMT mediates the IRE1-dependent effect. Moreover, comet assays quantify net DNA damage but do not distinguish altered damage formation from repair. Future studies incorporating MGMT depletion and rescue experiments, together with analyses of specific DNA repair-pathways and clonogenic-survival, would help clarify the functional contribution of MGMT to this phenotype. Second, the association of XBP1s with the MGMT promoter supports a potential transcriptional role but does not establish productive transcriptional regulation. Direct measurements of DNA methylation would be necessary to determine whether the DAC-associated response is specifically attributable to demethylation of the MGMT-promoter. Similarly, although the RNA-association and sequence analysis are consistent with RIDD-like regulation, definitive evidence of direct targeting would require RNA-stability and cleavage assays, rescue with RNase-defective IRE1 mutants, and mutational analysis of the candidate recognition element. The pooled ChIP-qPCR and CLIP-qPCR experiments provide evidence of molecular association but do not permit estimation of biological variability between independent experiments. Furthermore, construct-specific normalization and difference in expression levels warrants caution when comparing outcomes from IRE1 re-expression and XBP1s overexpression studies. Validation in additional cell models, together and with acute genetic or pharmacological perturbation of IRE1 signaling would further strengthen the mechanistic conclusions. Within these boundaries, the present study identifies a context-dependent relationship among IRE1 signaling, MGMT expression, and ETP-associated DNA damage. Collectively, the findings highlighting the importance of metabolic and stress-signaling states can influence the regulation of a DNA repair factor and the impact of IRE1 on genome maintenance.

## Materials and Methods

### Cell culture

MDA-MB-231 human triple-negative breast cancer cells were obtained from the American Type Culture Collection (ATCC; HTB-26, Manassas, VA, USA). Cells were maintained in glucose-free Dulbecco’s modified Eagle medium (DMEM; Gibco, Thermo Fisher Scientific, Waltham, MA, USA; cat. no. 11966025) supplemented with 10% fetal bovine serum (FBS; Dot Scientific, Burton, MI, USA) and D-glucose to a final concentration of 5.5 mM (Gibco Glucose Solution, Thermo Fisher Scientific; cat. no. A2494001). Cells were cultured at 37 °C in a humidified incubator with 5% CO₂. IRE1-deficient (IRE1^-/-^) MDA-MB-231 cells were generated by CRISPR/Cas9-mediated gene editing and validated as previously described (ref. 14). The same IRE1^-/-^ cell population was used in this study. IRE1 WT cells refer to parental MDA-MB-231 cells. Experiments were performed using cells between passages 8 to 25.

### Cell treatments and transfection

Cells were transfected with 10 nM ON-TARGETplus Human *XBP1* siRNA SMARTpool (Horizon Discovery; cat. no. L-009552-00-0005) or ON-TARGETplus Non-targeting Control Pool (Horizon Discovery; cat. no. D-001810-10-05) using Lipofectamine 3000 Transfection Reagent (Thermo Fisher Scientific; cat. no. L3000150) according to the manufacturer’s instructions. Cells were incubated for 48 h after siRNA transfection before chemical treatment.

For overexpression and re-expression experiments, MDA-MB-231 IRE1^-\-^ cells were transfected with 1,000 ng of empty vector or plasmids encoding human IRE1, XBP1s, or MGMT using Lipofectamine 3000 according to the manufacturer’s instructions. Cells were incubated for 24 h after plasmid transfection before chemical treatment. Equal total amounts of plasmid DNA were used across conditions.

For DAC (5-aza-2′-deoxycytidine; Sigma-Aldrich, A3656) experiments, MDA-MB-231 IRE1^-\-^ cells were transfected at approximately 50-60% confluency with 1,000 ng of empty vector or plasmids encoding human IRE1 or XBP1s, as described above. At 24 h after transfection, cells were passaged at a 1:4 split ratio. At 48 h after transfection, cells were treated with 100 nM DAC or volume-matched DMSO vehicle for 72 h. Treatment medium was replaced with fresh medium containing DAC or DMSO every 24 h. Cells were harvested at 120 h after transfection, corresponding to 72 h of DAC or vehicle treatment. The experimental schedule is provided in Supplementary Table S1.

Sodium palmitate (PA; Sigma-Aldrich; cat. no. P9767) was used at a final concentration of 0.1 mM. For PA+ETP experiments, cells were pretreated with PA for 8 h and subsequently co-treated with PA and 10 µM etoposide (ETP; Sigma-Aldrich; cat. no. E1383) for 16 h. ETP-only controls received 10 µM ETP for 16 h in the absence of PA. For pharmacologic experiments, WT MDA-MB-231 cells were treated with 30 µM 4μ8C (Sigma-Aldrich; cat. no. SML0949) or volume-matched DMSO vehicle for 3 h before, and during, a subsequent 3-h exposure to 10 µM ETP. Detailed cell-treatment and transfection procedures are provided in the Supplementary Methods.

### Comet assay

DNA damage was assessed using the Comet Assay Kit (3-well slides; Abcam, ab238544) according to the manufacturer’s instructions under alkaline electrophoresis conditions. Briefly, cells were harvested, resuspended in ice-cold PBS at 1×10^6^ cells/mL, mixed with Comet Agarose, and loaded onto the supplied comet slides. Slides were incubated in lysis buffer for 1 h at 4°C, followed by alkaline unwinding for 20 min at 4°C. Electrophoresis was performed in alkaline running buffer at 300 mA for 15 min. Slides were rinsed, fixed in 70% ethanol for 5 min, air-dried, and stained with Vista Green DNA Dye. Images were acquired using a Nikon ECLIPSE Ts2R inverted fluorescence microscope equipped with a 20x objective and controlled using NIS-Elements software (version 5.10; Nikon Instruments). Comet parameters were quantified using the OpenComet v1.3 ImageJ plugin. Tail moment was used as the primary indicator of DNA damage (ref. 17).

### Immunoblotting

Cells were lysed on ice for 15 min in RIPA Lysis and Extraction Buffer (Thermo Fisher Scientific, Waltham, MA, USA; cat. no. 89901) supplemented with 1× Halt Protease and Phosphatase Inhibitor Cocktail (Thermo Fisher Scientific, Waltham, MA, USA; cat. no. 78440). Lysates were clarified by centrifugation at 21,130×g (max) for 20 minutes at 4 °C, and protein concentrations were determined using the Pierce BCA Protein Assay Kit (Thermo Fisher Scientific, Waltham, MA, USA; cat. no. 23225) according to the manufacturer’s instructions. Equal amounts of protein (15 µg/lane) were separated on 4-15% or 8-16% Mini-PROTEAN TGX precast gels (Bio-Rad Laboratories, Hercules, CA, USA) and transferred to 0.2-µm PVDF membranes using the Trans-Blot Turbo Transfer System and Ready-to-Assemble Mini PVDF Transfer Kits (Bio-Rad Laboratories) according to the manufacturer’s instructions.

Membranes were blocked in EveryBlot Blocking Buffer (Bio-Rad Laboratories, Hercules, CA, USA) for 5 min at room temperature and incubated with primary antibodies diluted 1:1,000 in EveryBlot Blocking Buffer for 1 h at room temperature. Membranes were washed 5×5 minutes in TBST and incubated with HRP-conjugated secondary antibodies diluted 1:10,000 in the respective blocking buffer for 1 h at room temperature.

Chemiluminescent signals were developed using Clarity Western ECL Substrate (Bio-Rad Laboratories, Hercules, CA, USA; cat. no. 1705061) and acquired using a ChemiDoc MP Imaging System (Bio-Rad Laboratories). Band intensities were quantified using Image Lab software (version 6.1.0; Bio-Rad Laboratories) and normalized to vinculin or β-actin. Antibody details are provided in Supplementary Table S2.

### Analysis of mRNA levels by real-time quantitative PCR

Total RNA was isolated using the miRNeasy Tissue/Cells Advanced Mini Kit (QIAGEN, Hilden, Germany; cat. no. 217604) according to the manufacturer’s instructions. RNA concentration and purity were assessed by NanoDrop spectrophotometry (Thermo Fisher Scientific), and samples with A260/280 ratios of 1.8-2.1 were used for cDNA synthesis. cDNA was generated from 200 ng total RNA using the High-Capacity RNA-to-cDNA Kit (Applied Biosystems, Thermo Fisher Scientific; cat. no. 4387406) according to the manufacturer’s protocol. qRT-PCR was performed using PowerTrack SYBR Green Master Mix (Applied Biosystems, Thermo Fisher Scientific; cat. no. A46109) on a QuantStudio 5 Real-Time PCR System (Applied Biosystems, Thermo Fisher Scientific), following the manufacturer’s instructions. The primer sequences were as follows (5′-3′): MGMT, forward GCAATTAGCAGCCCTGGCA and reverse CACTCTGTGGCACGGGAT; and 18S rRNA, forward ATGGCCGTTCTTAGTTGGTG and reverse CGCTGAGCCAGTCAGTGTAG. The MGMT primer pair was also used for CLIP-qPCR. Reactions were performed in technical duplicate, and melt-curve analysis was used to confirm amplification specificity. Relative expression was calculated using the 2^−ΔΔCt^ method, normalized to 18S rRNA, and expressed relative to the indicated control condition.

### Cross-linking and chromatin immunoprecipitation followed by qPCR

Cross-linking immunoprecipitation followed by qPCR (CLIP-qPCR) was used to assess IRE1-associated MGMT RNA, whereas chromatin immunoprecipitation followed by qPCR (ChIP-qPCR) was used to assess XBP1s association with the MGMT promoter. For CLIP-qPCR, MDA-MB-231 WT cells transfected with XBP1-targeting or non-targeting siRNA were exposed to combined PA and ETP treatment as described above. Cells were UV-crosslinked at 254 nm with an energy dose of 150 mJ/cm² over 10 min. IRE1-associated RNA was immunoprecipitated using the anti-IRE1 antibody listed under immunoblotting, with IgG immunoprecipitates serving as negative controls. Recovered RNA was purified, reverse-transcribed, and analyzed using the MGMT transcript primers described above. MGMT transcript recovery was evaluated relative to IgG controls.

For ChIP-qPCR, MDA-MB-231 IRE1^-\-^ cells overexpressing XBP1s were treated with DAC or DMSO as described above. Following formaldehyde crosslinking and chromatin fragmentation, XBP1s-associated chromatin was immunoprecipitated using an anti-XBP1s antibody, with IgG and input controls included. Recovered DNA was analyzed using primers targeting the assayed *MGMT* promoter region. Enrichment was calculated as 2^−ΔCt2^, where ΔCt = Ct_IP_ - Ct_input_, using the input Ct values supplied in the service provider’s analysis.

Following crosslinking and sample collection, both assays were processed by Creative BioMart (Shirley, NY, USA). For each assay, material from three independent biological experiments was pooled for each condition, generating one pooled sample per condition that was measured in three technical qPCR replicates. Results were summarized descriptively as mean ± SEM of technical measurements, without inferential statistical testing between conditions. Detailed procedures and quality-control data are provided in the Supplementary Methods.

### *In silico* analysis of candidate IRE1 RNase recognition elements in MGMT mRNA

The human MGMT transcript sequence, MGMT-205 (ENST00000651593; gene ID: ENSG00000170430), was obtained from Ensembl release 112 using the GRCh38 human genome assembly. The source gene was located on chromosome 10 at positions 129,465,190– 129,770,983 on the positive strand. The transcript sequence was screened for the IRE1-associated RNA recognition motif, 5′-CUGCAG-3′, and the surrounding sequence context was evaluated for its potential to form an XBP1-like stem-loop structure. This sequence–structure combination has previously been associated with IRE1-mediated RNA recognition and cleavage. For each identified motif, a sequence window comprising the motif and 40-50 nucleotides upstream and downstream was extracted and analyzed using the RNAfold web server (ViennaRNA package; accessed July 20, 2026) under default parameters. Minimum-free-energy RNA secondary structures were predicted for each sequence window and examined for the position of the motif relative to the predicted loop and flanking paired stem. CUGCAG-containing regions located within the loop of a predicted stem-loop were considered candidate IRE1 recognition elements. The predicted MGMT RNA structure and candidate recognition motif are shown in Fig. 4d. This analysis was used to assess the structural plausibility of IRE1 recognition and was not considered evidence of direct RNA cleavage.

### Statistical analysis

Statistical analyses were performed using GraphPad Prism version 11 (GraphPad Software, Boston, MA, USA) and Microsoft Excel. Comet assay data were analyzed using two-sided Mann–Whitney U tests in GraphPad Prism 11. For western blot densitometry, qRT-PCR, and ChIP-qPCR analyses, fold-change or relative-enrichment values were log10-transformed before statistical testing. Comparisons between WT and IRE1-deficient cells in Fig. 1a-b were analyzed using two-sided Welch’s t-tests, which do not assume equal variances between groups. Comparisons in Fig. 2, Fig. 3, Fig. 4a-b, Fig. S1 and Fig. S2a-b were analyzed using two-sided paired t-tests, with samples paired within the same independent biological experiment. Data in Fig. 4c and Fig. S2c were analyzed using two-sided Welch’s t-tests. Both Welch’s t-tests and paired t-tests were performed in Microsoft Excel. Data are presented as mean ± SEM unless otherwise stated. All tests were two-sided, and *p* < 0.05 was considered statistically significant. The numbers of independent biological replicates and, where applicable, technical replicates are indicated in the corresponding figure legends.

## Declarations

### Generative AI use

This manuscript was prepared and edited with the assistance of Microsoft Copilot (Microsoft Corporation), accessed through Michigan State University’s enterprise platform. The tool was used to identify mechanical errors, improve clarity and concision, and suggest potentially relevant literature. All content, interpretations, and cited references were reviewed and verified by the authors. The authors take full responsibility for the accuracy, originality, and integrity of the manuscript.

### Ethical Approval

Ethical approval was not required for this study.

### Consent for Publication

Not applicable.

### Author Contributions

All authors contributed to methodology, conceptualization, data acquisition, investigation and manuscript reviewing. Kevin Chen wrote the original draft.

### Data Availability Statement

The data supporting the findings of this study are available from the corresponding author upon reasonable request.

