## Supplemental File for "Context-dependent regulation of MGMT expression by IRE1 signaling and its relationship to etoposide-associated DNA damage in MDA-MB-231 cells"

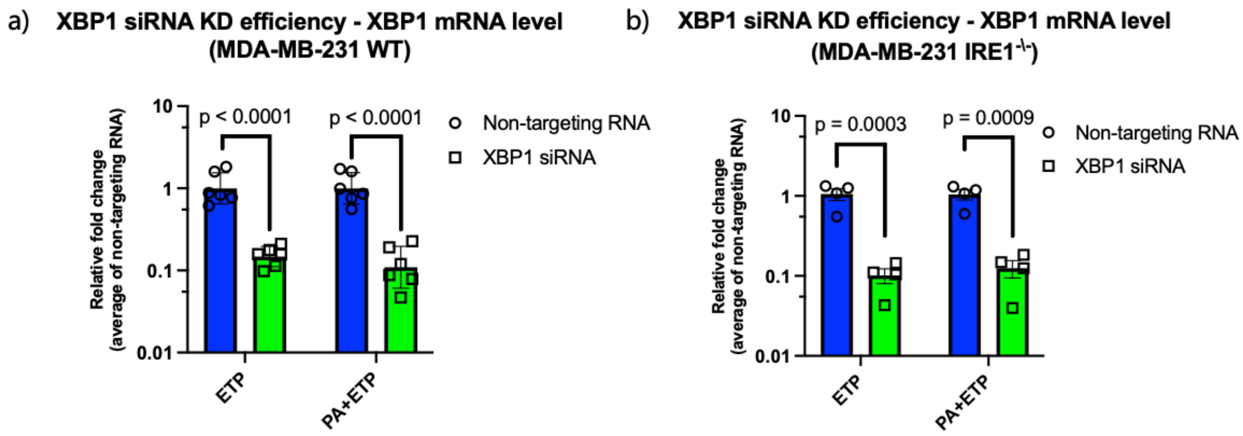


**Fig. S1. Validation of XBP1 knockdown in MDA-MB-231 WT and RE1$^{-/-}$ cells.** qRT-PCR analysis of XBP1 mRNA level in (a) WT and (b) RE1$^{-/-}$ cells transfected with XBP1-targeting siRNA (XBP1 KD) or non-targeting siRNA (NT) and treated with PA+ETP. Data are presented as mean ± SEM from n = 4 independent biological replicates.

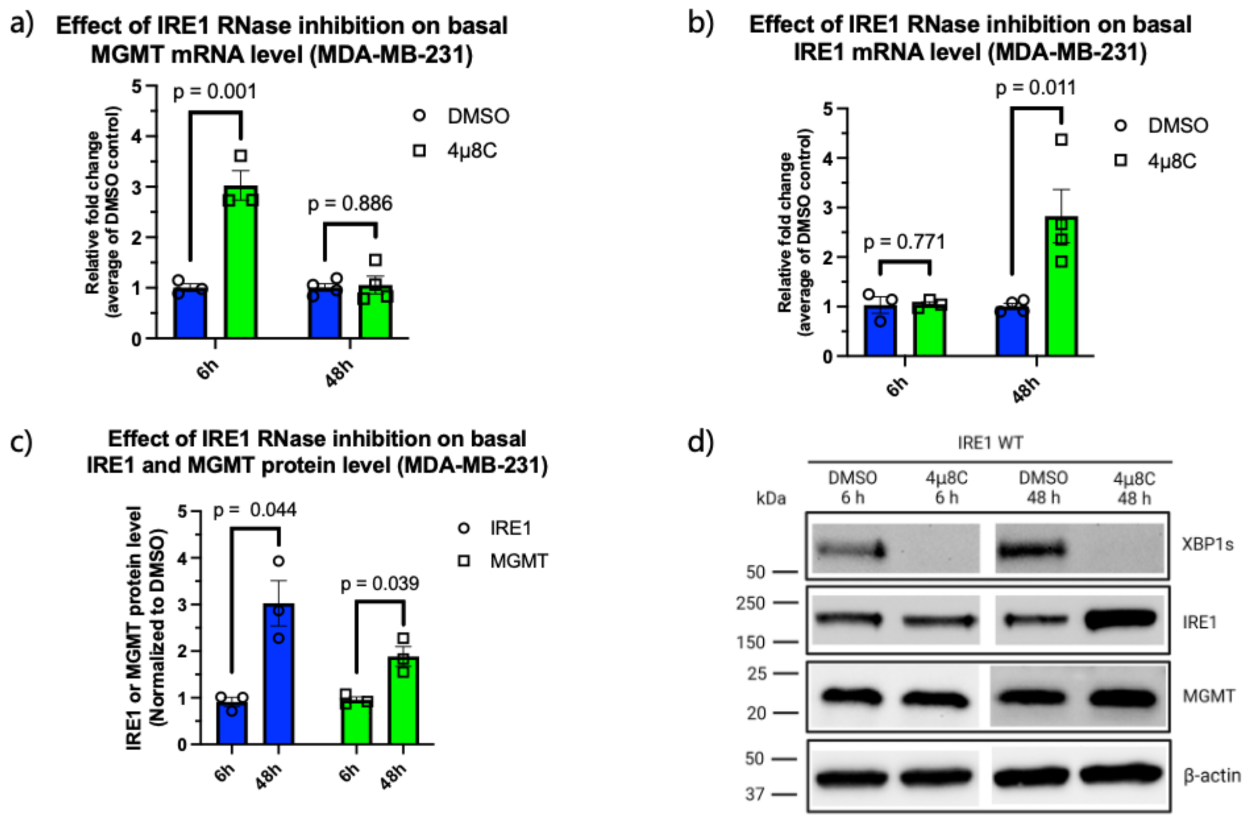


**Fig. S2. Time-dependent effects of IRE1 RNase inhibition on MGMT and IRE1 expression.** MDA-MB-231 cells were treated with 30 μM 4μ8C or DMSO vehicle for 6 or 48 h. For the 48 h treatment, 4μ8C were replenished at 24 h. (a, b) Relative MGMT (a) and IRE1 (b) mRNA levels measured by qRT-PCR and normalized to 18S rRNA. Compared with the corresponding DMSO controls, MGMT mRNA level was significantly increased at 6 h but not at 48 h, whereas IRE1 mRNA level was significantly increased at 48 h but not at 6 h. (c, d) Immunoblot analysis and quantification of IRE1 (c) and MGMT (d) protein levels, normalized to β-actin. Both proteins were significantly increased at 48 h, whereas neither showed a significant change at 6 h relative to the corresponding DMSO controls. Data are presented as mean ± SEM from n = 3 independent biological experiments.

| Day | Time after transfection | Procedure | Approximate confluency |
|---|---|---|---|
| 1 | 0 h | Transfect with 1,000 ng of empty vector, IRE1 plasmid, or XBP1s plasmid | 50–60% |
| 2 | 24 h | Passage cells at a 1:4 split ratio | – |
| 3 | 48 h | Initiate treatment with 100 nM DAC or volume-matched DMSO | 20–30% |
| 4 | 72 h | Replace treatment medium with fresh medium containing 100 nM DAC or DMSO | 50–60% |
| 5 | 96 h | Replace treatment medium with fresh medium containing 100 nM DAC or DMSO | 75–90% |
| 6 | 120 h | Harvest cells | 90–95% |

**Supplementary Table S1. Schedule for DAC treatment following IRE1 re-expression or XBP1s overexpression.** MDA-MB-231 IRE1$^{-\backslash-}$ cells were transfected with 1,000 ng of empty vector or plasmids encoding IRE1 or XBP1s. Cells were passaged at a 1:4 split ratio 24 h after transfection, and treatment with 100 nM DAC or volume-matched DMSO was initiated at 48 h. Treatment medium was refreshed every 24 h, and cells were harvested after 72 h of treatment, corresponding to 120 h after transfection. Confluencies indicate approximate values at each step.

| Target | Host/clonality | Vandor | Catalog no. | Expected molecular weight (kDa) |
|---|---|---|---|---|
| DNMT1 | Rabbit mAb | Abcam | ab188453 | ~183 |
| β-Actin | Rabbit mAb | Cell Signaling Technology | 8457 | 45 |
| IRE1 | Rabbit mAb | Cell Signaling Technology | 3294 | 130 |
| XBP1 | Rabbit mAb | Abcam | ab220783 | ~54 |
| MGMT | Rabbit mAb | Abcam | ab108630 | ~21 |
| Secondary | Goat polyclonal | Thermo Fisher Scientific | 31460 | - |

**Supplementary Table S2. Primary and secondary antibodies used for western blotting.** The table lists the target antigen, host species and clonality, vendor, catalog number, and expected molecular weight for each antibody used for immunoblot analysis. Antibody working dilutions are described in the Western blotting Methods section.

**Supplementary Methods**

Detailed cell-treatment and transfection procedures

MDA-MB-231 cells were seeded one day before transfection in 70-mm tissue-culture-treated dishes (GenClone; cat. no. 25-201) at $8\times10^5$ cells per dish for siRNA experiments or $1\times10^6$ cells per dish for plasmid-transfection experiments. Cells were transfected with 10 nM XBP1-targeting siRNA or non-targeting control siRNA using Lipofectamine 3000 Transfection Reagent (Thermo Fisher Scientific; cat. no. L3000150) according to the manufacturer's instructions. siRNA-transfected cells were incubated for 48 h before chemical treatment.

For siRNA-mediated *XBP1* knockdown, cells were transfected with 10 nM ON-TARGETplus Human *XBP1* siRNA SMARTpool (Horizon Discovery, Cambridge, UK; cat. no. L-009552-00-0005) or 10 nM ON-TARGETplus Non-targeting Control Pool (Horizon Discovery; cat. no. D-001810-10-05) using Lipofectamine 3000 Transfection Reagent (Thermo Fisher Scientific, Waltham, MA, USA; cat. no. L3000150) according to the manufacturer's instructions. Cells were incubated for 48 h after transfection before PA and/or ETP treatment.

For plasmid overexpression and IRE1 re-expression experiments, MDA-MB-231 IRE1$^{-\backslash-}$ cells were transfected with 1,000 ng plasmid DNA per 70-mm dish using Lipofectamine 3000 according to the manufacturer's instructions. Cells received a GenScript custom empty pcDNA3.1(+) vector (SC1317) or GenScript pcDNA3.1+/C-(K)-DYK expression plasmids encoding human IRE1/ERN1 (Clone ID OHu25357; RefSeq NM_001433.5), XBP1s (Clone ID OHu25513; RefSeq NM_001079539.1) or MGMT Clone ID OHu26767C; RefSeq NM_002412.4). The expression constructs contained C-terminal DYKDDDDK tags. Equal total amounts of plasmid DNA were used across conditions. Cells were incubated for 24 h after transfection before PA and/or ETP treatment.

Sodium palmitate (PA; Sigma-Aldrich, St. Louis, MO, USA; cat. no. P9767) was dissolved in water to prepare a 30 mM stock solution by heating until fully dissolved. Immediately before use, the PA stock was rapidly diluted into complete glucose-supplemented DMEM to a final concentration of 0.1 mM. For PA+ETP experiments, cells were pretreated with 0.1 mM PA for 8 h and then co-treated with 0.1 mM PA and 10 µM ETP (Sigma-Aldrich, St. Louis, MO, USA; cat. no. E1383) for an additional 16 h. ETP-only conditions received 10 µM ETP for 16 h in the absence of PA. Cells not treated with PA received complete culture medium without PA.

For IRE1 RNase-inhibition experiments, WT MDA-MB-231 cells were treated with 30 µM 4µ8C (Sigma-Aldrich, St. Louis, MO, USA; cat. no. SML0949) or volume-matched DMSO vehicle for 3 h before the addition of 10 µM ETP. Cells remained in 4µ8C or DMSO during a subsequent 3-h ETP exposure before processing for Comet assay.

CLIP-qPCR: UV crosslinking and sample preparation

MDA-MB-231 WT cells transfected with XBP1-targeting siRNA or non-targeting control siRNA were exposed to combined PA and ETP treatment as described in the main Methods. Following treatment, the medium was removed, and cells were washed twice with ice-cold PBS. Cells in 10-cm dishes were covered with 1 mL of ice-cold PBS and UV-crosslinked with the dish lids removed at 254 nm with an energy dose of 150 mJ/cm² over 10 min. An additional 9 mL of ice-cold PBS was added, and cells were collected by scraping and centrifugation at 1,200 rpm for 3 min at 4°C. Pellets were washed once with 10 mL of ice-cold PBS, collected by centrifugation, snap-frozen in liquid nitrogen, and stored at -80°C before subsequent processing by Creative BioMart.

CLIP-qPCR: Immunoprecipitation and RNA analysis

UV-crosslinked cell pellets were washed with pre-cooled PBS and resuspended in CLIP lysis buffer containing protease inhibitors. Lysates were treated with RNase and DNase and sonicated for five cycles of 30 s on and 30 s off. Following centrifugation at 12,000 rpm for 10 min at 4°C, clarified supernatants were retained for immunoprecipitation. An aliquot of 50 µL was reserved as input. The remaining lysates were incubated with anti-IRE1 antibody or control IgG overnight at 4°C with rotation. Antibody-associated complexes were captured using magnetic beads for 4 h at 4°C and washed with CLIP binding buffer.

Complexes were eluted at 37°C for 5 min with agitation at 1,000 rpm. Eluates were treated with proteinase K at 56°C for 15 min followed by 72°C for 15 min. RNA was recovered using TRIzol and a micro RNA extraction kit, reverse-transcribed, and analyzed by qPCR. The MGMT transcript primers were forward 5′-GCAATTAGCAGCCCTGGCA-3′ and reverse 5′-CACTCTGTGGCACGGGAT-3′. Each 20-µL qPCR reaction contained 10 µL of Power SYBR Green Master Mix, 0.5 µL each of 10 µM forward and reverse primers, 1 µL of cDNA, and 8 µL of water. Cycling conditions were 95°C for 1 min followed by 40 cycles of 95°C for 15 s and 63°C for 25 s, with fluorescence acquisition, followed by melt-curve analysis from 55°C to 95°C. MGMT transcript recovery in anti-IRE1 immunoprecipitates was evaluated relative to IgG controls.

ChIP-qPCR: Crosslinking and sample preparation

MDA-MB-231 IRE1$^{-/-}$ cells overexpressing XBP1s were treated with DAC or DMSO as described in the main Methods. Cells were fixed with 1% formaldehyde for 10 min at room temperature with gentle agitation. Fixation was quenched by adding 2.5 M glycine at a 1:20 ratio (500 µL per 10 mL of fixation solution), followed by incubation for 5 min at room temperature with gentle agitation. Following quenching, cells were washed twice with PBS, collected by scraping, and centrifuged at 1,200 rpm for 3 min at 4°C. Pellets were resuspended in 1 mL of PBS, transferred to microcentrifuge tubes, and centrifuged again under the same conditions. Cell pellets were snap-frozen in liquid nitrogen and stored at -80°C before subsequent processing by Creative BioMart.

ChIP-qPCR: Immunoprecipitation and DNA analysis

Crosslinked cells were resuspended in ChIP lysis buffer and sonicated using 30-s on/30-s off cycles at the provider-reported 99% power setting for a reported total duration of 40 min. Lysates were centrifuged at 12,000 rpm for 10 min at 4°C, and the supernatants were retained. An aliquot of 50 μL was reserved as input, and 450 μL of lysate was used for anti-XBP1s immunoprecipitation. Samples were incubated with antibody overnight at 4°C with rotation, followed by incubation with 50 μL of Protein G magnetic beads for 4 h at 4°C. IgG immunoprecipitates served as negative controls. Beads were washed four times before elution.

Complexes were eluted at 37°C for 5 min with agitation at 1,000 rpm. Following proteinase K digestion and crosslink reversal according to the service provider's procedure, DNA was purified using a micro DNA purification kit. The assayed MGMT promoter region was amplified using forward primer 5′-CTTTCAGGACCACTCGGGC-3′ and reverse primer 5′-ATGGCCCGTACCTTTTCCTG-3′. Each 50-μL reaction contained 25 μL of SYBR Premix Ex Taq II, 2 μL each of 10 μM forward and reverse primers, 2 μL of DNA template, and 19 μL of nuclease-free water. Cycling conditions were 95°C for 30 s followed by 40 cycles of 95°C for 5 s and 60°C for 30 s. Amplification was evaluated using melt curves and agarose gel electrophoresis. Input-normalized enrichment was calculated as described in the main Materials and Methods.
